# An Entropy-Based Approach to Model Selection with Application to Single-Cell Time-Stamped Snapshot Data

**DOI:** 10.1101/2025.01.03.631247

**Authors:** William CL Stewart, Ciriyam Jayaprakash, Jayajit Das

## Abstract

Recent single-cell experiments that measure copy numbers of over 40 proteins in individual cells at different time points [time-stamped snapshot (TSS) data] exhibit cell-to-cell variability. Because the same cells cannot be tracked over time, TSS data provide key information about the time-evolution of protein abundances that could yield mechanisms that underlie signaling kinetics. We recently developed a generalized method of moments (GMM) based approach that estimates parameters of mechanistic models using TSS data. However, when multiple mechanistic models potentially explain the same TSS data, selecting the best model (i.e., model selection) is often challenging. Popular approaches like Kullback-Leibler divergence and Akaike’s Information Criterion are difficult to implement because the distribution that gave rise to the “noisy” data is only known numerically and approximately. To perform model selection in this situation, we introduce an entropy-based approach that incorporates our GMM based parameter estimation and commonly used estimators in kernel density estimation. Using simulated TSS data, we show that our approach can select the “ground truth” from a set of competing mechanistic models. Furthermore, we use a bootstrap procedure to compute model selection probabilities, which can be useful when measuring the relative support of a candidate model.

## 1 Introduction

Ordinary differential equations (ODEs) are commonly used to model the subcellular dynamics of proteins and mRNA [14, 15, 18]. Usually, ODEs describe the deterministic dynamics of average concentrations, which facilitates the estimation of reaction rates (*aka* model parameters) from experimental data. This is often an important step towards building mechanistic biological models. But the task of estimating model parameters from experimental data is challenging for a variety of reasons ([2, 17, 25]), especially when the number of distinct proteins measured in experiments is smaller than the number of model parameters. Recent developments in single-cell experimental techniques for the longitudinal measurements of transcripts and proteins in an individual cell (e.g., RNA-seq [20] which can simultaneously measure over a thousand different RNA sequences, and CyTOF [16, 24] which can simultaneously measure more than thirty different protein species) appear to alleviate this problem. Since individual cells are not tracked across time in these experiments, the measurements generate a large collection of time-stamped snapshot (TSS) data. Another challenge stems from the cell-to-cell differences in the copy number of a protein (or abundance), which contains variation present at the pre-stimulus state (*aka* extrinsic noise), and variation arising from the stochasticity of biochemical reactions over time (*aka* intrinsic noise) [7, 26]. When the observed protein abundances are large, extrinsic noise is known to play a significant role [8] and single-cell protein signaling kinetics can be well approximated by ODEs. By comparing the distribution of protein abundances observed at time *t* to the predictions obtained from an ODE model that evolves protein abundances seen at an earlier time (e.g. *t* = 0 *aka* initial conditions), we can estimate the parameters of the candidate model [13]. In this paper, we propose an entropy-based approach to address the larger question of model selection for systems that can be described by deterministic dynamical models (e.g. sets of ODEs) with randomness arising from initial conditions. Specifically, for the models considered here, we show that our crossentropy [4] approach can find the *best* ODE model from a set of competing candidate models, where the best model neither overfits nor underfits the available data.

The primary goal of model selection is to find the best model relative to some defensible criterion, and two attractive criteria are cross-entropy and Kullback-Leibler (KL) divergence [19]. The latter is a non-negative number that measures the “distance” between two probability distributions [13]. Usually, the distribution that gave rise to the protein abundances observed at time *t* (denoted *f*) is considered to be the “ground truth”, while the other distribution is most often a candidate distribution (or model) that is presumed to be “close” to *f*. A defining feature of KL divergence is that it is zero if and only if the candidate model and *f* are the same. Typically, the best candidate model will strike a balance between underfitting (i.e. under-estimating systematic variation and over-estimating noise) and overfitting (i.e. over-estimating systematic variation and under-estimating noise). Since KL divergence is a difference in expectations taken with respect to *f* (i.e. cross-entropy minus entropy). and since entropy depends only on *f*, it suffices (for the purpose of model selection) to find the model with the smallest cross-entropy. Given a finite set of candidate models, the model with the smallest cross-entropy is also the model with the smallest KL divergence, which makes it the best approximating model to *f*.

There are however two very important challenges when performing model selection from TSS data. First, the multivariate probability density of the observed protein abundances at time *t* (denoted symbolically as *f*, and in words as the “ground truth”) is rarely known. Second, while we can estimate the parameters of each candidate model, we cannot evaluate the likelihood of any model. Fortunately, we can deal with the first challenge by minimizing crossentropy (instead of KL divergence), as cross-entropy only requires *realizations* from *f*, not complete knowledge of *f*. As for the second challenge (which cannot be avoided), we tackle it head on by estimating each candidate model (see Methods and Appendix B for more details) from the time evolution of initial conditions (see Data Description and Appendix A for more details).

The remaining sections of this paper are organized as follows. In the first subsection of Methods, we give a mathematical descriptions of KL divergence and cross-entropy, and we briefly explain how parameters of a candidate model are estimated. Then, in the second subsection of Methods, we outline our model selection approach leaving technical details (such as the estimation of the Gaussian copula, marginal densities, and marginal cumulative distribution functions) to Appendices A and B. Furthermore, in the second subsection of Methods, we briefly describe a complementary model selection approach that applies Akaike Information Criterion [1] corrected for small samples (AICc) [3] to differences between the mean protein abundances observed at time *t* and the mean protein abundances predicted at time *t*. In Data Description, we describe how synthetic TSS data are generated, and we describe the time-evolution of initial conditions using different candidate models. Finally, we demonstrate the utility of our model selection approach in Results, and give some interesting insights and suggestions for further improvements in Discussion.

## 2. Methods

### 2.1 Kullback-Leibler and Cross-Entropy

Consider the following example of TSS data: protein abundances for *n* distinct proteins observed at time zero in a single cell (denoted *x*[0]), and protein abundances for the same set of distinct proteins observed at time *t* in a different cell (denoted *y*[*t*]). We assume that *x*[0] ∼ *G, y*[*t*] ∼ *F* with density *f* (*y*[*t*]) and *y*[0] (which can never be observed) has the same distribution as *x*[0]. Furthermore, in accordance with most real-world applications, we assume that *G, F*, and *f* are almost never known. Now, for *i* ∈ (1, 2, 3) let *d*_*i*_ be a set of coupled nonlinear ODEs that governs the deterministic dynamics of single-cell protein abundances over time, where each *d*_*i*_ has reaction rates (see Fig. 1) that depend on freely varying parameters *θ*_*i*_. We evolve initial conditions *x*[0] to time *t* using ODE model *d*_*i*_ to arrive at predicted abundances *x*_*i*_[*t*] which have distribution *H*_*i*_ and density *h*_*i*_ referred to interchangeably as the *i*^th^ candidate model. For each candidate model, we assume there exists a unique parameter vector 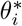 that minimizes

**Figure 1.**
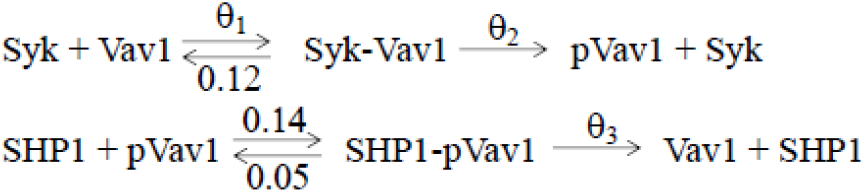
Minimal NK cell Signaling Model: Shows the biochemical reactions for the six protein species interaction model, Syk, Vav1, Syk-Vav1, SHP1, and SHP1-pVav1. The ODE model describes the deterministic mass action kinetics corresponding to the reactions shown here. To ensure that all reaction rates are identifiable, three were fixed at 0.12, 0.14, and 0.05, respectively.

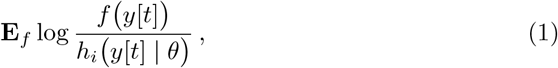

where the expectation in expression (1) is the Kullback-Leilber (KL) divergence between *f* (*y*[*t*]) and candidate model *h*_*i*_(*y*[*t*] | *θ*). Note that, when the *i*^th^ candidate model and the “ground truth” are the same, *h*_*i*_ = *f* and the KL divergence is zero. Since we have restricted attention to a single point in time, we suppress the dependence of *x*_*i*_[*t*] and *y*[*t*] on *t* hereafter.

KL divergence can also be expressed as the difference between cross-entropy and entropy

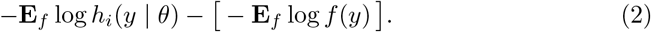

So, when 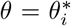 expression (2) is the KL divergence between *f* and *h*_*i*_. Since entropy does not depend on *h*_*i*_, it suffices (for the purpose of model selection) to find the candidate model that minimizes cross-entropy

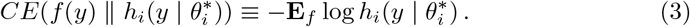

However, because *f* is rarely known, the cross-entropy in Eq.(3) cannot be computed. Instead, we must approximate the cross-entropy by averaging over 𝒩 independent realizations of *f*. To compute the approximation we also need 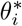 and *h*_*i*_, but unfortunately, both are rarely known as well. In the case of 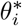, we can replace it with the generalized method of moments (GMM) estimate of *θ* (denoted 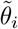). Briefly as in [13], we define 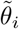 to be the value of *θ* that minimizes the weighted sum of squared differences between the first and second moments of *y*, and the first and second moments of *x*_*i*_ [11]. Note that for most over-determined systems, 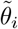 is consistent for 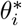 [10]. In the next subsection, we will describe how *h*_*i*_ can be estimated from initial conditions *x*[0], coupled ODEs *d*_*i*_, and corresponding estimated reaction rates 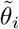. Our estimate of *h*_*i*_ is denoted 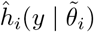.

### 2.2 Computing Approximate Cross-Entropy

Single-cell experiments such as CyTOF typically measure protein abundances across thousands of cells, and as such, TSS data often contain a wealth of information for estimating marginal densities and marginal distribution functions. Consequently, we decided to leverage Sklar’s theorem [23] to estimate the multivariate density 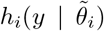 from its marginal densities, marginal distribution functions, and a copula describing the dependence between the abundances of *n* distinct proteins in a single cell (see Appendix B for more details). Here, we implement a Gaussian copula, which is computationally fast, mathematically convenient, and (at least in the case of TSS data) quite accurate (see Appendix C).

Another benefit of working with large samples, is that split-sample techniques allow us to avoid the bias that typically arises when parameter estimation and model selection are performed on the same dataset [1]. Specifically, we use 20% of the available TSS data for parameter estimation (i.e. to compute 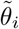), and 80% for model selection (i.e. to compute 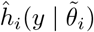. Now, the cross-entropy that we want (see Eq. 3), can be approximated by

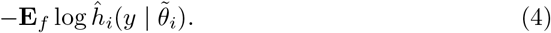

and the expression in (4) can be estimated from the corresponding sample average (denoted ACE):

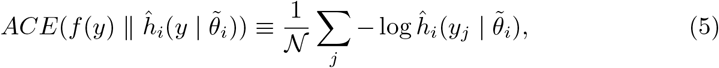

where 𝒩 independent cells at times 0 and *t* are used to compute 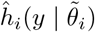 and ACE, respectively.

Because our proposed model selection approach is based on an approximation, it is useful to have (1) independent confirmation that the selected model *actually* minimizes KL divergence, and (2) some measure of relative support for the model that ACE selects compared to the other models that could have been chosen. In particular, by modeling the mean of *y* minus the mean of *x*_*i*_ as a multivariate normal random vector with expectation zero, and by assuming a variance-covariance matrix with off-diagonal elements equal to zero, we can approximate AICc [3] for the *i*^th^ candidate model when the number of distinct proteins *n* is strictly larger than the maximum number of freely varying parameters (which is the case for all of the candidate models that we consider here). Note that ACE has no such limitation. When the candidate model that minimizes ACE also minimizes approximate AICc, one has (to some degree) additional assurance that the selected model minimizes cross-entropy [see Eq.(3)], and therefore minimizes KL divergence [see Eq. (2)]. Furthermore, by bootstrapping the observed TSS data, we can estimate the *model selection* probabilities for each candidate model (i.e. the probability that model *h*_*i*_ is selected). Typically, model selection probabilities will help users quantify and interpret the level of relative support for the selected model in ways that are comparable to, or better than, differences in AIC or AICc [5].

## 3 Data Description

In most real-world applications that model the time-dependent kinetics of protein abundances, the “ground truth” (denoted *f*) is rarely known. However, because we are primarily concerned with improving model selection (i.e. increasing accuracy, increasing usability, and developing additional measures of relative support), we mimic observed data *y* by simulating TSS data (without intrinsic noise) from each of three candidate models. In particular, when we pretend that *f* = *h*_1_ (which depends on one freely varying parameter), we refer to this scenario as “ground truth” SMALL. Similarly, when *f* = *h*_2_, and *f* = *h*_3_, we refer to these scenarios as “ground truth” MEDIUM and “ground truth” LARGE, respectively. Note that, for candidate model *h*_1_, *θ*_2_ = 9 \**θ*_1_ and *θ*_3_ = 2 * *θ*_1_ leaving only *θ*_1_ to vary freely. For candidate model *h*_2_, *θ*_2_ = 9 * *θ*_1_ and parameters *θ*_1_ and *θ*_3_ vary freely. For candidate model *h*_3_, parameters *θ*_1_, *θ*_2_, and *θ*_3_ are all allowed to vary freely. Hence, *h*_1_ is nested within *h*_2_, and *h*_2_ is nested within *h*_3_. Furthermore, despite never having a closed-form expression for the “ground truth” in any scenario, we always know which candidate model the “ground truth” is in every scenario. Of course, when implementing ACE (or AICc), we pretend that *f* is unknown.

To generate TSS data, we begin by simulating uncorrelated initial conditions from a multivariate lognormal distribution with parameters: *µ* = (5.25, 7.60, 5.25, 7.60, 5.25, 5.25), and *σ*^2^ = (0.15, 0.06, 0.15, 0.06, 0.15, 0.15). Now, let’s consider simulated data corresponding to “ground truth” LARGE; this scenario has three freely varying parameters. We take half of the initial conditions (discussed immediately above) and we evolve them to time *t* using coupled ODEs *d*_3_ and estimated parameter vector 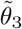. Then, for the remaining initial conditions, we evolve them using *d*_1_, *d*_2_, and *d*_3_ together with their corresponding estimated parameters 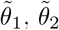, and 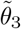 to carry out model selection.

## 4 Results

We show that our model selection approach works, and that when approximate AICc can be computed, it selects the same candidate model as ACE (Table 1).

**Table 1.** Model Selection Based on Simulated TSS Data. For each “Ground Truth” nonlinear model (LARGE, MEDIUM, and SMALL), we generated synthetic time-stamped snapshot (TSS) data for 6 proteins across 8,000 cells at time *t* = 1.5*s*. Columns 2 and 3 show the minimum ACE and the second smallest ACE, respectively. Columns 4 and 5 show the minimum AICc and the second smallest AICc, respectively. The minimization is taken over all three candidate models. The selected candidate model (CM) is shown in brackets, with *CM*_3_ ≡ *h*_3_, *CM*_2_ ≡ *h*_2_, and *CM*_1_ ≡ *h*_1_. For both ACE and AICc, the correct candidate model is always selected.

| “Ground Truth”<br>Scenario | Minimum<br>ACE | 2nd Smallest<br>ACE | Minimum<br>AICc | 2nd Smallest<br>AICc |
| --- | --- | --- | --- | --- |
| LARGE | 23.8 [$CM_3$] | 24.2 [$CM_2$] | 33 [$CM_3$] | 54 [$CM_2$] |
| MEDIUM | 22.5 [$CM_2$] | 23.6 [$CM_3$] | 20 [$CM_2$] | 32 [$CM_3$] |
| SMALL | 23.5 [$CM_1$] | 23.6 [$CM_2$] | 10 [$CM_1$] | 21 [$CM_2$] |

Now, we repeated our model selection approach for each of 1000 bootstrap resamples of the simulated TSS data, and computed the probability that the correct candidate model was actually selected. Recall that, in our simulation, the “ground truth” model is always one of the candidate models under consideration. For the “ground truth” LARGE scenario, ACE selected the correct model 100% of the time. Similarly, for the “ground truth” MEDIUM scenario, the minimum ACE selected the correct model 95% of the time, and for the “ground truth” SMALL scenario, the accuracy of ACE was 76% (see Table 2).

**Table 2.**
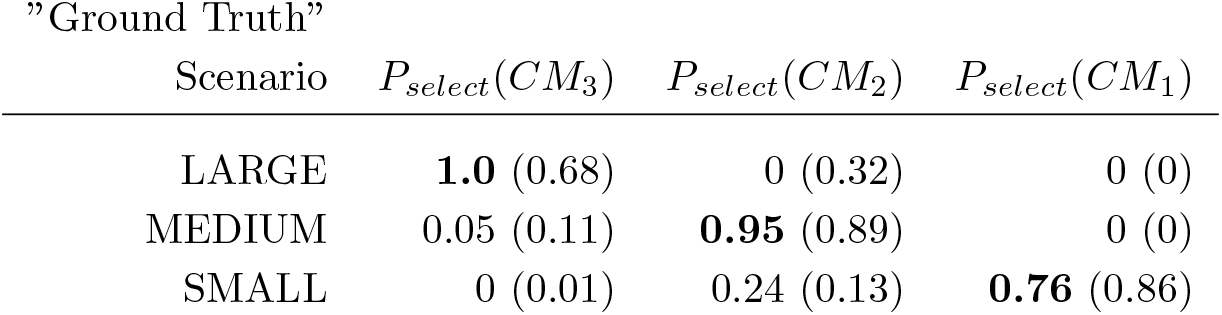
Accuracy of ACE Model Selection: For each “Ground Truth” simulated TSS data set, 1000 bootstrap resamples were generated. Here, we show the probability of selecting candidate models: *CM*_3_ ≡ *h*_3_, *CM*_2_ ≡ *h*_2_, and *CM*_1_ ≡ *h*_1_ using ACE when the “Ground Truth” scenario is LARGE, MEDIUM, or SMALL, respectively. Corresponding probabilities for AICc are given in parentheses.

## 5 Discussion

### 5.1 New Tools for Model Selection with Single-Cell Data

Mechanistic models based on ODEs describing subcellular kinetics of proteins are widely used in computational biology for gleaning mechanisms and generating predictions. It is common to have multiple candidate mechanistic models that can be set up to probe different hypotheses describing the same biological phenomena, and an important task in model development is to rank order the candidate models according to their ability to describe the measured data. The availability of high dimensional single cell data sets allow for estimation of model parameters using mean values and higher order moments of the measured data, however, rank ordering models calibrated against these datasets may not be straightforward using standard model selection tools such as AICc. Here we propose a model selection approach based on cross-entropy for ODE based models that are calibrated against means and higher order moments of the measured data. We show as “proof-of-concept” that our proposed approach successfully rank orders a set of ODE models against synthetic single cell datasets.

### 5.2 ARE Complements Approximate AICc

As shown in Fig.(2), approximate AICc is virtually independent of ACE and concordance between ACE and approximate AICc appears to provide additional support for the selected candidate model. So, any improvement in approximate AICc would like only benefit ACE. One potential area for improvement might be a recalibration of the penalties used in approximate AICc. In particular, it’s not completely clear what the penalty should be for approximate AICc, as the parameter estimation is based on thousands of cells, but the model selection is based on differences in only six means.

**Figure 2.**
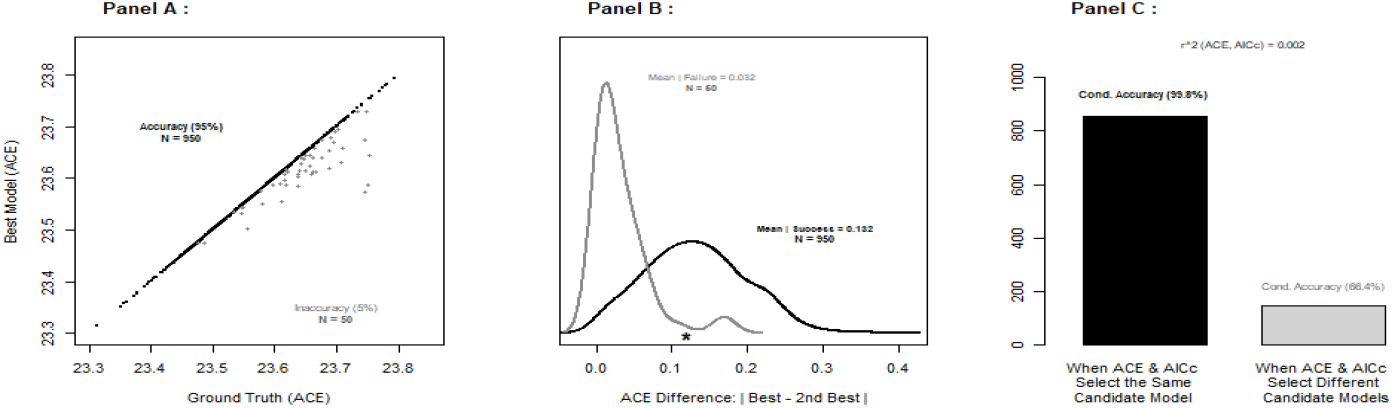
A Detailed Examination of the “Ground Truth” MEDIUM Scenario: Based on 1000 bootstrap resamples, Panel A shows the overall accuracy (black dots) and inaccuracy (grey dots) of ACE for the “ground truth” MEDIUM scenario. Panel B shows the distribution of absolute differences between the smallest and second smallest ACE scores when ACE selects the correct model (black) or an incorrect model (grey). The asterisk is the absolute difference between the ACE of *h*_2_ and *h*_3_ for data simulated with the “ground truth” MEDIUM scenario. Panel C shows how concordance with AICc provides additional evidence that the model selected by ACE is correct.

A requirement for approximate AICc is that the number of distinct proteins must be larger than the number of freely varying parameters, otherwise the denominator of the approximate AICc statistic will be zero or negative. ACE does not have this limitation. Also, as approximate AICc assumes independence between abundances observed at time *t*, there is a distinct possibility that its performance may decline in the presence of a non-trivial correlation structure. Since ACE estimates the full multivariate distribution, it should be able to handle whatever correleations exist in the data.

For the MEDIUM and LARGE “ground truth” scenarios, ACE outperforms approximate AICc, but for the SMALL “ground truth” scenario approximate AICc does better than ACE. This probably happens because the means tend to carry the lion’s share of the information in the “ground truth” SMALL scenario, so there’s very little benefit to estimating the other candidate models (which contain information about higher order moments and cross moments). However, when the higher order moments and cross moments begin to matter, as is likely the case with the more complex “ground truth” scenarios, the benefit of estimating the candidate models with 2 and 3 freely varying parameters is likely much greater.

### 5.3 Limitations and Future Directions

Presently, our ACE approach to model selection based on TSS data has three main limitations: (1) it is not designed to handle intrinsic noise, (2) there is considerable latitude in terms of multivariate density estimation that we have only scratched the surface of, and (3) there may be more efficient ways to incorporate split-sample techniques.

Here, we tailored our analysis to simulated TSS data, but for other applications users may want to include higher order moments and/or different copulas or kernel density estimators [12, 21]. Furthermore, we have extrinsic noise only, no intrinsic noise (or at least) intrinsic noise is assumed to be small. Extending our ACE approach to include *both* extrinsic and intrinsic noise may be possible for relatively short evolution times and/or for networks with a relatively small number of interacting proteins.

When researchers are unwilling or unable to specify the full model, the likelihood is not known and model selection is often challenging. However, when consistent estimators of the model parameters exist, and when the sample size is large (in the number of independent observations at time *t*, and in the number of samples with initial conditions), we propose a split-sample entropy-based approach to model selection that allows users to find the best approximating model to the “ground truth”. Furthermore, our approach is quite flexible with respect to (1) parameter estimation (e.g. choosing which moments to use… first moments only, first and second moments, etc.), and (2) multivariate density estimation (e.g. choosing a “good” kernel density estimator and/or copula for each candidate model).

## 6 Appendix A: Evolving Protein Abundances with ODEs

We consider a simplified model of biochemical reactions that describe early time signaling kinetics in mouse Natural Killer (NK) cells stimulated by ligands cognate to activating CD16 and inhibitory Ly49A receptors. We explicitly model the kinase Syk bound to activating receptor-ligand complex and the phosphatase SHP1 bound to inhibitory receptor-ligand complex. The enzyme Syk phosphorylates a key signaling protein Vav1 where phosphorylated Vav1 (pVav1) induces activation of NK cells. The enzyme SHP1 dephosphorylates pVav1 and thus mediates inhibition of NK cell activation. In our model, Syk and SHP1 react with the substrate protein Vav1 and pVav1, respectively, following the reactions shown in (Fig. 1). Activating and inhibitory NK cell receptors are not included explicitly in this model; instead, Syk and SHP1 represent Syk and SHP1 proteins bound to the activating and inhibitory receptors, respectively. The reactions in the model are similar to that of the zero-order ultrasensitivity model proposed by Goldbeter and Koshland ([9]). The abundances for the protein species, Syk, Vav1, Syk-Vav1, SHP1, SHP1-pVav1 and pVav1 evolve in time with a set of nonlinear ODEs describing mass-action kinetics for the above reactions.

To describe the signaling kinetics of protein abundances *y*, which evolve in time through *r* biochemical signaling reactions (see Figure 1) according to a set of coupled ODEs we use

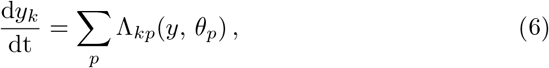

where *k* ∈ (1, …, *n*) indexes the distinct proteins observed in a single cell at time *t*. Here, Λ_*kp*_(*y, θ*_*p*_) is the propensity of the *p*^th^ reaction affecting the abundance of the *k*^th^ protein, and *θ*_*p*_ is the rate constant for the *p*^th^ biochemical signaling reaction. The ODEs allow for both linear and nonlinear dynamics, and the parameter values for all simulated data are taken (in part) from the parameters used in [6]. For a given model of time-evolution, the ODEs are solved numerically using a Runge-Kutta Cash-Karp nonlinear solver in C++ at time *t* = 1.5 *s* to generate synthetic data *y* at time *t* and to evolve initial conditions *x*[0] for each candidate model: *h*_1_, *h*_2_, and *h*_3_.

## 7 Appendix B: Estimating Candidate Models

Let *i* ∈ (1, 2, 3) index the candidate models, and let *ĥ* _*i*_(*y*) be the estimate of the *i*^th^ candidate model *h*_*i*_(*y*). Now, let *j* ∈ (1, …, *n*) index the number of distinct proteins observed in a cell. To compute *ĥ* _*i*_(*y*) we will need: (1) estimates of the marginal cumulative distribution functions *H*_*ij*_, (2) estimates of their corresponding marginal densities *h*_*ij*_, and (3) an estimate of the correlation structure between protein abundances.

Let *x*_*i*_ be the evolution of initial condition *x*[0] to time *t* based on a set of coupled ODEs that depend on parameters 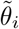. The *n* components of *x*_*i*_ = (*x*_*i*1_, …, *x*_*in*_) are continuous representations of protein abundances for *n* distinct proteins. Now, since *x*_*ij*_ is predicted in *m* single cells indexed by *k* = 1, …, *m* and since *m* is assumed to be large (say *>* 1000), the marginal cumulative distribution function of *x*_*ij*_ is well estimated by:

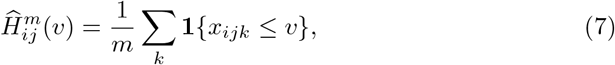

where *x*_*ijk*_ is the predicted abundance for the *j*^th^ protein in the *k*^th^ cell, 0 *< v <* ∞, and **1** is the indicator function that takes the value 1 when the event *{x*_*ijk*_ ≤ *v}* is true. By the strong law of large numbers,

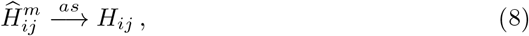

as *m* → ∞. Furthermore, to estimate the corresponding marginal density functions *h*_*ij*_, we use the default kernel density estimation procedure [22] in the statistical software package R (version 4.2.1). The estimate of *h*_*ij*_ is denoted *ĥ*_*ij*_

To estimate the correlation structure, for each *j* we apply a *double-transform* to *x*_*ij*_, so that the resulting random vector *z*_*i*_ has a multivariate normal distribution with mean zero and variance-covariance Σ. Specifically, we define *u*_*ijk*_ ≡ *Ĥ*_*ij*_(*x*_*ijk*_) and *z*_*ijk*_ ≡ Φ^−1^(*u*_*ijk*_), so that (*z*_*i*1*k*_, … *z*_*ink*_) ∼ *MV N* (0, Σ) for each *k*. Then, our estimate of the correlation structure (denoted 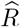) is simply the standard estimate of Σ appropriately rescaled. Finally, our estimate of the *i*^th^ candidate model evaluated at *y* is

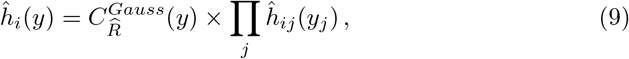

where *y* ≡ (*y*_1_, …, *y*_*n*_) represents protein abundances observed at time *t* in a single cell, and

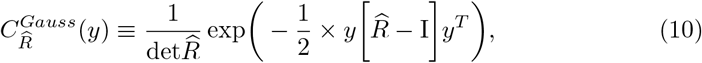

with I being the [*n × n*] identity matrix. Hence, our estimate of the *i*^th^ log-likelihood of 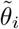 based on *y* is log *ĥ*_*i*_(*y*), and now, the computation of ACE in Eq. (5) is straightforward.

## 8 Appendix C: Accuracy of Gaussian Copula

To assess the accuracy of the Gaussian copula, we compared the exact log-likelihood of simulated initial conditions where the distribution of protein abundance for the *i*^th^ distinct protein is known to be multivariate log normal (*µ*_*i*_, 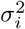) and uncorrelated with the abundances of other proteins (see Data Description for more details). The log-likelihood differences shown in Table (3) suggests that the error in our log-likelihood calculations for ACE is only slightly greater than 1%.

**Table 3.** Accuracy of the Gaussian Copula: 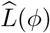 We generated initial conditions for 6 distinct proteins across 3,000 and 6,000 cells, respectively using a multivariate log-normal distribution indexed by *ϕ* ≡ (*µ, σ*^2^). The Gaussian copula over-estimated the exact log-likelihood by slightly more than 0.5% when *N* = 3000; and it over-estimated the exact log-likelihood by slightly less than 1% when *N* = 6000.

| Sample Size | $\log L(\phi)$ | $\log \hat{L}(\phi)$ |
| --- | --- | --- |
| 3,000 | -5859.07 | -5821.99 |
| 6,000 | -11718.14 | -11611.02 |

## 9 Acknowledgements

This work was supported by NIH grant R01AI146581 to J.D. and by GIG Statistical Consulting LLC, which provided computational resources for the simulations and data analyses. We thank John Wu for developing the C++ code to solve the non-linear ODEs.

## 11 Author Contribution

Conceptualization, W.C.L.S; methodology, W.C.L.S, C.J., J.D.; software, W.C.L.S; validation, W.C.L.S; formal analysis, W.C.L.S; investigation, W.C.L.S; resources, W.C.L.S; data curation, W.C.L.S; writing—original draft preparation, W.C.L.S, C.J., J.D.; writing—review and editing, W.C.L.S, C.J., J.D.; visualization, W.C.L.S, C.J., J.D.; supervision, W.C.L.S, C.J., J.D.; project administration, W.C.L.S, C.J., J.D.; funding acquisition, J.D. All authors have read and agreed to the published version of the manuscript.

## References

[1] H Akaike. “A new look at the statistical model identification”. IEEE Transactions on Automatic Control 19.6 (1974), pp. 716–723. doi: 10.1109/TAC.1974.1100705.

[2] Maksat Ashyraliyev, Yves Fomekong-Nanfack, Jaap A Kaandorp, and Joke G Blom. “Systems biology: parameter estimation for biochemical models”. The FEBS journal 276.4 (2009), pp. 886–902.

[3] J. E. Cavanaugh. “Unifying the derivations of the Akaike and corrected Akaike information criteria”. Statistics & Probability Letters 31.2 (1997), pp. 201–208. doi: 10.1016/s0167-7152(96)00128-9.

[4] T. M. Cover and J. A. Thomas. Elements of Information Theory, 2nd Edition. Wiley, 2006.

[5] A Dajles and J Cavanaugh. “Bootstrap Approximation of Model Selection Probabilities for Multimodel Inference Frameworks”. Entropy 26.599 (2024), pp. 1–17. doi: 10.3390/e26070599.

[6] Jayajit Das. “Activation or tolerance of natural killer cells is modulated by ligand quality in a nonmonotonic manner”. Biophysical journal 99.7 (2010), pp. 2028–2037.

[7] Jayajit Das and Ciriyam Jayaprakash. Systems immunology: An introduction to modeling methods for scientists. CRC Press, 2018.

[8] Ofer Feinerman, Joël Veiga, Jeffrey R Dorfman, Ronald N Germain, and Grégoire Altan-Bonnet. “Variability and robustness in T cell activation from regulated heterogeneity in protein levels”. Science 321.5892 (2008), pp. 1081–1084.

[9] Albert Goldbeter and Daniel E Koshland. “An amplified sensitivity arising from covalent modification in biological systems”. Proceedings of the National Academy of Sciences 78.11 (1981), pp. 6840–6844.

[10] Alastair R. Hall and Atsushi Inoue. “The large sample behaviour of the generalized method of moments estimator in misspecified models”. Journal of Econometrics 114.2 (2003), pp. 361–394. issn: 0304-4076. doi: 10.1016/S0304-4076(03)00089-7.

[11] Lars Peter Hansen. “Large sample properties of generalized method of moments estimators”. Econometrica 50 (1982), pp. 1029–1054.

[12] T. Hastie, R. Tibshirani, and J. H. Friedman. The Elements of Statistical Learning : Data Mining, Inference, and Prediction : with 200 full-color illustrations. New York: Springer, 2001.

[13] Wu J, Stewart WCL, Jayaprakash C, and Das J. “BioNetGMMFit: estimating parameters of a BioNetGen model from time-stamped snapshots of single cells”. npj Systems Biology and Applications 46.9 (2023).

[14] Gioele La Manno, Ruslan Soldatov, Amit Zeisel, Emelie Braun, Hannah Hochgerner, Viktor Petukhov, Katja Lidschreiber, Maria E Kastriti, Peter Lönnerberg, Alessandro Furlan, et al. “RNA velocity of single cells”. Nature 560.7719 (2018), pp. 494–498.

[15] Pavel Loskot, Komlan Atitey, and Lyudmila Mihaylova. “Comprehensive review of models and methods for inferences in bio-chemical reaction networks”. Frontiers in Genetics 10 (2019), p. 549.

[16] Sayak Mukherjee, Helle Jensen, William Stewart, David Stewart, William C Ray, Shih-Yu Chen, Garry P Nolan, Lewis L Lanier, and Jayajit Das. “In silico modeling identifies CD45 as a regulator of IL-2 synergy in the NKG2D-mediated activation of immature human NK cells”. Science signaling 10.485 (2017).

[17] Andreas Raue, Clemens Kreutz, Thomas Maiwald, Julie Bachmann, Marcel Schilling, Ursula Klingmüller, and Jens Timmer. “Structural and practical identifiability analysis of partially observed dynamical models by exploiting the profile likelihood”. Bioinformatics 25.15 (2009), pp. 1923–1929.

[18] Jennifer A Rohrs, Pin Wang, and Stacey D Finley. “Understanding the dynamics of T-cell activation in health and disease through the lens of computational modeling”. JCO clinical cancer informatics 3 (2019), pp. 1–8.

[19] Kullback S and Leibler RA. “On information and sufficiency”. Annals of Mathematical Statistics 22.1 (1951), pp. 79–86.

[20] Antoine-Emmanuel Saliba, Alexander J Westermann, Stanislaw A Gorski, and Jörg Vogel. “Single-cell RNA-seq: advances and future challenges”. Nucleic acids research 42.14 (2014), pp. 8845–8860.

[21] S. J. Sheather and M. C. Jones. “A reliable data-based bandwidth selection method for kernel density estimation”. Journal of the Royal Statistical Society, Series B 53.3 (1991), pp. 683–690. doi: 10.1111/j.2517-6161.1991.tb01857.x.JSTOR2345597.

[22] B. W. Silverman. Density Estimation for Statistics and Data Analysis. London: Chapman and Hall, 1986.

[23] Abel Sklar. “Fonctions de répartition à n dimensions et leurs marges”. 8 (1959), pp. 229–231.

[24] Matthew H Spitzer and Garry P Nolan. “Mass cytometry: single cells, many features”. Cell 165.4 (2016), pp. 780–791.

[25] William CL Stewart. “The fundamentals of statistical data analysis”. In: Systems Immunology. CRC Press, 2018, pp. 41–50.

[26] Peter S Swain, Michael B Elowitz, and Eric D Siggia. “Intrinsic and extrinsic contributions to stochasticity in gene expression”. Proceedings of the National Academy of Sciences 99.20 (2002), pp. 12795–12800.

